# Controlled delivery of iNOS antagonist, 1400W, for synergistic breast cancer therapy

**DOI:** 10.64898/2026.05.28.728138

**Authors:** Houman Alimoradi, Marjan Abri Aghdam, Anita Fallah, Christine Delporte

## Abstract

Triple-negative breast cancer (TNBC) is an aggressive subtype of breast cancer that lacks effective targeted therapies and is frequently associated with chemotherapy resistance and immunosuppression. Inducible nitric oxide synthase (iNOS) is overexpressed in breast cancer and has been strongly correlated with poor clinical outcomes, owing to its role in promoting tumor progression, invasiveness, and resistance to therapy. Although highly selective iNOS inhibitors such as N-(3-(Aminomethyl)benzyl)acetamidine (1400W) exhibit considerable therapeutic promise, their clinical translation has been hindered by unfavorable pharmacokinetic properties. To overcome these limitations, we developed a pH-responsive nanoscale formulation based on Schiff base conjugation between 1400W and oxidized PEGylated alginate (OPA), in combination with ionic interactions. The resulting nanoparticles (NPs) exhibited efficient release of 1400W under acidic conditions and effectively suppressed nitric oxide (NO) production in lipopolysaccharide (LPS)-stimulated RAW264.7 macrophages. While the NPs alone did not induce significant cytotoxicity, they synergistically enhanced the anticancer efficacy of paclitaxel (PTX) in MDA-MB-231 TNBC cells, significantly inhibiting cell viability and migration. In addition, the NP–PTX combination markedly reduced endothelial tube formation in HUVECs, compared to PTX alone indicating potentiation of the anti-angiogenic activity of PTX. In conclusion, the pH-responsive NPs enables effective modulation of NO signaling and enhances the therapeutic activity of PTX in TNBC cells. These findings support the potential of iNOS-targeted nanomedicine as an adjuvant strategy for TNBC treatment and warrant further investigation using *in vivo* models to evaluate pharmacokinetics, tumor accumulation, and antitumor efficacy of the NPs.

## Introduction

Triple-negative breast cancer (TNBC) accounts for approximately 15–20% of all breast cancers and is associated with higher rates of metastasis, recurrence, and mortality than other subtypes, resulting in poor overall survival. TNBC is characterized by the absence of estrogen receptor, progesterone receptor, and human epidermal growth factor receptor 2 expression, which limits the availability of effective molecular targets for therapy [1, 2]. Consequently, treatment primarily relies on surgery, radiotherapy, and conventional chemotherapy to control tumor growth and reduce metastatic spread [1, 2]. However, systemic intravenous chemotherapy results in nonspecific drug distribution throughout the body, leading to significant toxicity while often failing to achieve adequate drug concentrations at tumor sites. Despite ongoing advances, treatment resistance and metastasis remain the leading causes of mortality in patients with TNBC. As a result, standard chemotherapy produces response rates of only approximately 20%, offering limited improvement in long-term survival [3].

Nitric oxide (NO) is a highly versatile mediator involved in immune defense, vascular regulation, neurotransmission, and metabolism. Nitric oxide synthase (NOS) catalyzes the conversion of L-arginine to NO and citrulline. NOS exists in three isoforms, endothelial (eNOS), neuronal (nNOS), and inducible (iNOS), which share a common catalytic mechanism but differ in their regulation and cellular distribution [4]. The biological effects of NO depend on multiple factors, including its concentration, duration of exposure, cellular redox state, and cell type. NO can either maintain physiological homeostasis or, under certain conditions, contribute to tissue injury. This dual role is particularly evident in cancer. Low concentrations of NO promote angiogenesis and tumor perfusion, thereby supporting tumor growth and metastasis, whereas high concentrations of NO induce DNA damage, inhibit mitochondrial respiration, and activate apoptotic pathways, resulting in antitumor effects [4, 5].

Elevated iNOS expression has been found in breast cancer [6, 7] as well as in various other cancers, including colon [8], lung, glioblastoma and melanoma lung [9], glioblastoma [10] and melanoma [11]. Preclinical and clinical studies have shown an association between elevated iNOS expression and poor prognosis, as well as aggressiveness in breast cancer patients [12–14]. In the tumor microenvironment (TME), iNOS is overexpressed in tumor-associated macrophages (TAMs) and cancer cells, creating a feedforward loop of inflammation and leading to therapeutic resistance [15]. Consequently, selectively inhibiting iNOS has emerged as a strategy to suppress tumor-promoting pathways while preserving the homeostatic functions of constitutive NOS isoforms [16]. A few ongoing clinical trials are investigating the therapeutic potential of iNOS inhibition for cancer treatment, including the use of NG-monomethyl-L-arginine (L-NMMA) in combination with taxane-based chemotherapy in patients with breast cancer [17]. Despite these promising outcomes, concerns remain regarding the off-target effects of NOS inhibitors, particularly hypertension and pulmonary symptoms [17, 18].

An acetamidine derivative 1400W (N-(3-(aminomethyl)benzyl) acetamidine) is a selective, tight-binding, and potent inhibitor of human iNOS [19]. It suppresses proliferation, migration, and invasion of iNOS-expressing tumor cells and induces apoptosis and cell-cycle arrest in several cancer models, including TNBC [6, 7], colorectal and pancreatic [20] cancers [8], and intrahepatic cholangiocarcinoma [21]. In mouse xenografts, it limits the growth of established tumors and reduces metastatic burden, effects commonly associated with decreased angiogenesis, inflammation, and cancer stem cell self-renewal. In addition, combining 1400W with standard chemotherapeutics such as 5-fluorouracil or PTX further enhances antiproliferative and antimetastatic responses, supporting its potential as an adjuvant strategy in future cancer therapy [22, 23].

However, clinical translation is constrained by a short plasma half-life, limited tumor accumulation, and dose-limiting systemic toxicity [10, 24]. Most iNOS antagonists are cationic, which leads to nonspecific interactions with negatively charged cell surfaces, poor pharmacokinetic behavior, and nonselective tissue distribution. As a result, higher doses are often required, further compromising their selectivity.

Nanoformulation is an emerging strategy to address these limitations by prolonging circulation time and improving tumor targeting of small molecules. This advantage is partly driven by the enhanced permeability and retention (EPR) effect [25–28], which promotes preferential accumulation in tumor tissue over normal tissue. To date, however, no delivery system has been developed to enhance the delivery of 1400W.

In this study, we developed a nanosized formulation for controlled delivery of 1400W based on Schiff base interactions between the aldehyde groups of oxidized PEGylated alginate (OPA) and 1400W. This approach yielded stable, pH-responsive nanoparticles that enable controlled release of 1400W and synergistically enhance the cytotoxicity of PTX.

## Materials and methods

Unless specified otherwise, chemicals used in this work were obtained from TCI-Belgium (Zwijndrecht, Belgium) or Sigma-Aldrich, (St. Louis, MI, USA). All solvents were purchased from Thermo Fisher Scientific (Waltham, MA, USA) or Merck (Carlsbad, CA, USA) and employed directly without further purification. NMR spectra were acquired on a 400 MHz spectrometer (JEOL, Tokyo, Japan) and processed using MestReNova software (version 15, CIREM licensed). Reagents for cell culture were provided by Thermo Fisher Scientific. FTIR spectra were acquired with an Alpha II spectrometer (Bruker Optics GmbH & Co. KG, Ettlingen, Germany) using attenuated total reflectance (ATR) mode in the 4000–500 cm□^1^ range, at a resolution of 1 cm□^1^, averaging 32 scans per sample. Zeta potential measurements were performed using a Zetasizer Ultra (Malvern Instruments Ltd., Malvern, UK). UV/Vis absorbance was measured using a LAMBDA™ 25/35 Series UV/Vis spectrophotometer (LAMBDA™ 25/35 Series UV/Vis spectrophotometer, PerkinElmer, Springfield, IL, USA) or Spark® Multimode Microplate Reader (Tecan, Männedorf, Switzerland). Flow cytometry analysis was performed using a FACScan flow cytometer (Becton Dickinson, NJ, USA), and the data were analyzed with FlowJo software (FlowJo, Ashland, OR, USA). Microscopy was carried out using LSM900 confocal microscope equipped with Airyscan (Carl Zeiss, Oberkochen, Germany). Transmission electron microscopy (TEM) was carried out at the Center for Microscopy and Molecular Imaging (CMMI, Université Libre de Bruxelles, Belgium) using Tecnai 10 100 kV instrument (Philips Electronics International B.V., Eindhoven, Netherlands).

### Synthesis of oxidized OPA

The carboxyl groups of sodium alginate (SA, 1 g) were activated with EDC (1 g; 1-ethyl-3-(3-dimethylaminopropyl) carbodiimide) and NHS (1 g; N-hydroxysuccinimide) in 100 mL of distilled water, followed by reaction with polyethylene-glycol (PEG)-amine (2 g, 2KDa). The mixture was maintained at 25 °C for 24 h. The product was precipitated by dropwise addition into 200 mL of acetone and centrifuged at 91g This purification step was repeated three times by redissolving in distilled water and reprecipitating with acetone. The final precipitate was washed with 10 mL acetone and dried using a rotary evaporator.

Subsequently, 2 g of the obtained PEGylated alginate was dissolved in 100 mL distilled water and reacted with NaIO□ (2 g) at 25 °C for 24 h to induce oxidation. Ethylene glycol (3 mL) was then added to quench unreacted NaIO□. Acetone (four times the reaction volume ~ 400 mL) and NaCl (1 g) were added to precipitate the OPA. The precipitate was transferred into a dialysis bag (molecular weight cutoff (MWCO) of 3.5kDa) and dialyzed against deionized water for 24 h.

### Determination of OPA oxidation degree

The oxidation degree (OD) of OPA was determined by UV–Vis spectroscopy based on the consumption of sodium periodate (NaIO□) before the addition of ethylene glycol [29]. Briefly, 0.1 mL of the reaction mixture was diluted to 25 mL with distilled water. An aliquot (400 µL) of this solution was mixed with 1.50 mL of 10% (w/v) potassium iodide (KI) and 0.5% (w/v) soluble starch solutions. The absorbance was recorded immediately at 290 nm using a UV/Vis spectrophotometer (PerkinElmer). The OD was calculated using the following equation: *OD= Ni-Nr/N0×100*, where: Ni is initial moles of NaIO□, Nr is residual moles of NaIO□ after the reaction, No is initial moles of uronic acid units in alginate.

### Formation of NPs from 1400W and OPA

A vigorously stirred solution of OPA (1 mg/mL) in 50 mL 10 mM phosphate-buffered saline (PBS, pH 7.4) was prepared, and 150 mg (0.6 mmol) of 1400W·2HCl in 10 mL of 20% (v/v) THF/water was added dropwise. The reaction was allowed to proceed at room temperature for 1 h. The mixture was then dialyzed against distilled water using a 3.5 kDa membrane for 48 h, followed by lyophilization to obtain the NPs.

The size, polydispersity index (PDI), and surface charge of the synthesized NPs were determined using DLS. Measurements were conducted at 25°C in phosphate-buffered saline (1X PBS, pH 7.4).

### Quantification of 1400W Loading and Release Profile

To assess the *in vitro* release profile of 1400W from NPs, 5 mL of NP solution (2 mg/mL) was loaded into dialysis bags (3.5 kDa MWCO) and submerged in 50 mL PBS pH 7.4. The release of 1400W from NPs was quantified using a ninhydrin-based colorimetric assay [30]. Briefly, 1 mL aliquots of the dialysate were transferred to glass tubes, followed by the sequential addition of 100 μL potassium cyanide (KCN, 3 mM) and 100 μL ninhydrin solution (4% in phenol). The tubes were sealed, vortexed, and incubated in a preheated water bath at 97°C for 10 min. After cooling to room temperature, the solutions were transferred into 2 mL volumetric flasks, and the volume was adjusted to 2 mL using ethanol. The absorbance of 100 μL of the final solution was measured at 570 nm using UV–Vis spectroscopy.

To quantify the loading of 1400W, NP solutions (1 mg/mL in distilled water) were adjusted to pH 12 using 1M NaOH and stirred for one hour. The solution was then extracted with ethyl acetate, and the organic layers were dried and redissolved in phenol. The concentration of 1400W was measured using a ninhydrin assay via spectrophotometry at 570 nm.

### Cell culture

The human triple-negative breast cancer (TNBC) line MDA-MB-231, murine macrophages RAW264.7 (ATCC), and human umbilical vein endothelial cells (HUVECs, ATCC, passages 6– 8) were from the American Type Culture Collection (ATCC) (Manassas, VA, USA). MDA-MB-231 and RAW264.7 cells were maintained in DMEM supplemented with 10% fetal bovine serum (FBS), 100 U/mL penicillin and 100 µg/mL streptomycin. HUVECs were cultured in endothelial growth medium (ATCC CRL-1730™) according to the supplier’s instructions, containing growth factors including human EGF, VEGF, hFGF-B, and R3-IGF-1, with additional 10% FBS, 100 U/mL penicillin and 100 µg/mL streptomycin. All cultures were grown as monolayers in a humidified incubator at 37□°C with 5% CO□.

### Quantification of Nitric Oxide (NO) in RAW264.7 Cells

RAW 264.7 murine macrophages (400,000 cells per well) were seeded into 24-well plates and incubated at 37°C in a humidified atmosphere with 5% CO□ for 24 h. The cells were then treated with lipopolysaccharide (LPS, 1 μg/mL), 1400W-loaded NPs and free 1400W for another 24 h. Following incubation, the cells were exposed to 4 μM 4,5-diaminofluorescein-2 diacetate (DAF-2Da), a fluorescent probe for NO detection, for 30 min at 37°C. The medium was then removed and replaced with fresh media, followed by staining with 1 μg/mL DAPI for 10 min in the dark. After washing, cells were visualized using fluorescence confocal microscopy as previously described [31]. For quantification of fluorescence intensity, 100,000 cells were seeded in 24-well plates, allowed to adhere overnight, and treated under the same conditions used for fluorescence imaging. After 24 h, cells were washed with PBS, trypsinized, resuspended in fresh medium, and stained with 4 µM DAF-2Da following the same protocol as for fluorescence imaging. the fluorescence intensity of stained cells was measured using flow cytometry, and the data were processed using FlowJo software (FlowJo, Ashland, OR, USA).

### Cell viability measurement

RAW 264.7 murine macrophages or MDA-MB-231 cells were seeded in 96-well plates at 7,000 cells per well and incubated for 24 h to allow attachment and growth. Cells were then incubated without or with different concentrations of the test compounds (PTX, NPs and their combinations) for 24 h. After treatment, the medium was replaced with 100 µL of MTT solution (0.4 mg/mL in fresh cell culture medium). Following a 3h incubation, the medium was removed, and the formed formazan crystals were solubilized in 100 µL of DMSO. Absorbance was read at 540 nm using a Victor X2 plate reader (PerkinElmer) to quantify cell viability.

### Calcein/propidium iodide staining assay

MDA-MB-231 cells were seeded in flat-bottom black 96-well plates at a density of 10,000 cells per well and allowed to adhere overnight under standard culture conditions. Following treatment of the cells with 1 µM PTX and/or 20 µM NPs, cells were stained with calcein (2 µM) and propidium iodide (PI; 1 μg/mL) according to the manufacturer’s instructions to assess cell viability. DAPI (1 μg/mL) was used for nuclear counterstaining. Fluorescent images were acquired using a fluorescent confocal microscope to visualize live (calcein-positive, in green) and dead (PI-positive, in red) cells.

### Scratch cell migration assay

A scratch assay was performed to evaluate inflammation-induced cell migration. 1×10^5^ MDA-MB-231 cells were seeded in 24 well-pates and allowed to grow for 24h. Confluent cells cultured in a 24-well plate were scratched using a sterile 200□µL pipette tip to create a uniform gap. Detached debris was removed by gently rinsing with PBS. Cells were then treated with 1 µM PTX and/or 20 µM NPs in serum-free medium and incubated for 24 h. Images of the scratch area were captured at 0 and 24 h using the microscope. Scratch width was measured using ImageJ software, and migration was calculated as the percentage of scratch closure over time.

### HUVEC tube formation assay

Matrigel was thawed on ice and carefully distributed into the wells of a 96-well plate (50μL/well), then allowed to solidify for 1 h at 37°C. HUVECs (10,000 cells per well) were subsequently seeded in culture medium supplemented with 5% fetal bovine serum and 100 U/mL100 U/mL penicillin and 100 µg/mL streptomycin. Cells were stimulated with VEGF (50 ng/mL) either alone or together with the NPs (40μM), PTX (1 μM), or a combination of the two (PTX+NPs). Following a 24 h incubation at 37°C, capillary-like network formation was visualized by light microscopy. Quantitative assessment of angiogenic parameters was performed using the Angiogenesis Analyzer plugin in Fiji (ImageJ).

### Statistical analysis

Statistical analyses were performed using a t-test or one-way ANOVA with Tukey’s post hoc test, Kruskal-Wallis test with Dann’s multiple comparison test in GraphPad Prism (v9.1.0), as appropriate. Gaussian distribution of data was verified by Kolmogorov-Smirnov test. Data are shown as mean ± SD and a p value < 0.05 was considered statistically significant. Combination effects (synergism and antagonism) were evaluated by the Chou–Talalay method using CompuSyn 1.01 (ComboSyn, Inc., Paramus, NJ, USA). Combination index (CI) values < 1, = 1, and > 1 indicated synergistic, additive, and antagonistic interactions, respectively.

## Results and Discussion

### Synthesis and characterization of the OPA

The preparation of NPs from 1400W was done through the interactions between amine groups of 1400W and aldehyde of oxidized SA (Figure 1a). As PEGylation of NPs surface has been a well-established process to prolong their circulation half-life by avoiding immune recognition and clearance [32], SA was initially PEGylated using EDC/NHS through the amide coupling of amine groups of a 2kDa PEG-NH2 and carboxylic groups of SA (Figure 1a). After purification, PEGylate SA was reacted with NaIO_4_ for different incubation periods to assess the oxidation degree of PEG-SA estimated to 28.3 and 46.1% after 5 and 24h incubation with 50% NaIO_4_ respectively.

**Figure 1.**
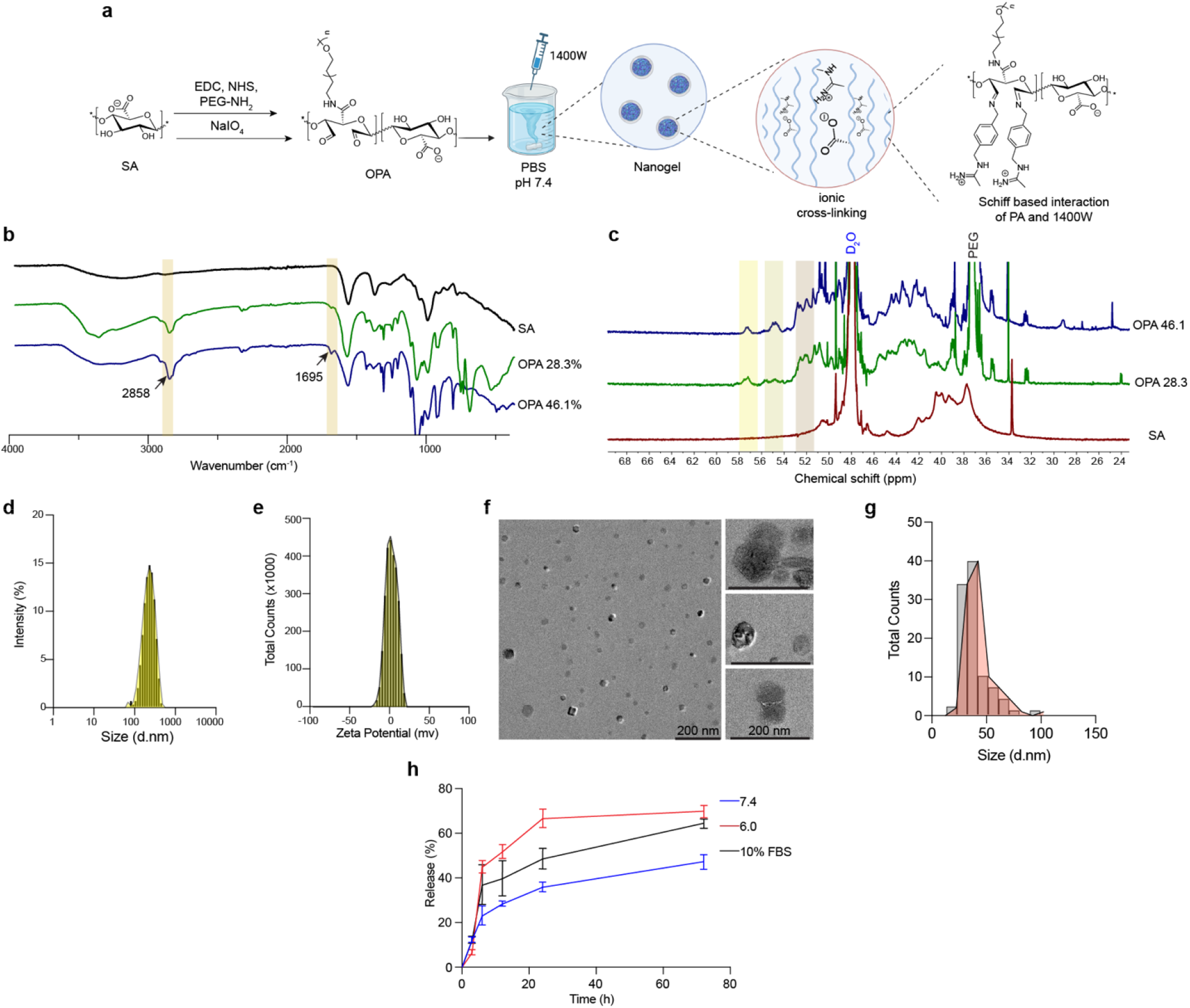
Synthesis and characterization of NPs made of 1400W and OPA. **a.** Schematic representation of the chemical synthesis of OPA and assembly of 1400W-loaded NPs. **b**. FT-IR of SA and OPA at oxidation degrees of 28.3% and 46.1%. **c**. ^1^H NMR of SA and OPA at oxidation degrees of 28.3% and 46.1%. **d**. Representative histogram of NP size distribution measured by DLS using a 1 mg/mL NP solution in distilled water. **e**. Zeta potential of NPs measured in 10 mM NaCl solution. **f**. Representative TEM images of the NPs. **g**. NP diameter obtained from TEM analysis. **h**. Release of 1400W profile of the NPs under different conditions (PBS pH 6.0 and 7.4 and PBS (pH 7.4) +10% FBS) over 72 h. Data are expressed as mean ± SD (n=3).

Following oxidation, the presence of aldehyde groups in OPA was assessed using FT-IR spectroscopy. The FT-IR spectrum of unmodified SA (Figure 1b) displays a broad peak around 3200 cm□^1^ corresponding to O–H stretching, a weak peak at ~2930 cm□^1^ related to aliphatic C– H stretching. The strong peaks observed at 1608 cm□^1^ and 1419 cm□^1^ are attributed to the asymmetric and symmetric stretching vibrations of carboxylate groups, which are well-documented as features of alginate [33, 34]. The peak at round 1100 cm□^1^ attributed to C-O stretching notably increased in OPA and this mainly could be due to PEG groups of OPA. The characteristic alginate peaks remained visible post-oxidation, however peaks at 1608 cm□^1^ (C=O stretching) slightly shifted towards higher wavenumbers and at high DO (46.1 %) a small peak at 1695 cm□^1^ was observed which could be due to some of the aldehyde carbonyls in the structure (Figure 1a). Notably, the intensity of C-H and O-H stretching peaks increased after oxidation, potentially due to elimination of intramolecular interactions and cleavage of glycosidic bonds and further increase of hydroxyl groups [34, 35].

The ^1^HNMR spectrum of OPA (Figure 1c) showed a strong peak at 3.6 ppm which belongs PEG. In addition, new peaks in the range of 5.1 ppm to 5.7 ppm (absent in the SA spectrum, particularly peaks at 5.3 ppm, 5.5 ppm, and 5.7 ppm) were observed which corresponded to hemiacetalic proton formed from aldehyde and neighboring hydroxyl groups [36–39].

### Preparation of NPs from 1400W and OPA

The interaction between 1400W and OPA leads to the formation of nanosized particles, likely in a nanogel format. Different ratios of OPA and 1400W were investigated for NP synthesis (Figure 1a). Among the tested formulations, particles formed from OPA with 28.3% degree of oxidation (DO) and an excess of 1400W yielded NPs with an average hydrodynamic size of 202 ± 26 nm (Figure 1d), a slightly negative surface charge (−2.9 ± 5.4 mV, Figure 1e), and a low PDI (0.2). The primary amine group of 1400W reacts with the aldehyde group of OPA to form a Schiff base. In addition, ionic interactions between positively charged acetimidate groups and negatively charged carboxylate groups further contribute to NP formation and result in a nearly neutral surface charge. These NPs exhibited a loading ratio of 15.9% (W/W) for 1400W. TEM analysis revealed spherical NPs with significantly smaller sizes (35.5 ± 12 nm) compared to those measured by DLS (202 ± 26 nm) (Figure 1g). This discrepancy is likely attributable to NP hydration in solution, which affects the hydrodynamic diameter measured by DLS, whereas TEM reflects the dehydrated state.

The NPs demonstrated good stability under physiological conditions (pH 7.4 in PBS) and showed sustained release of 1400W over 72 h. Notably, the release rate was significantly increased in the presence of 10% fetal FBS, likely due to ionic interactions between FBS components and the NPs that compromise NP stability. Furthermore, the NPs exhibited a pH-responsive release profile. Within the first 12 h, 1400W release at pH 6 was 1.8-fold higher (51.6%) than that observed at pH 7.4 (28.3%), and a similar trend persisted up to 72 h (Figure 1h).

Various characteristics of the TME have been exploited for targeted drug delivery in cancer therapy [40]. Among these, pH-responsive systems have been extensively investigated [41], owing to the acidic nature of tumors (pH 6.5–6.8) compared with normal tissues (pH 7.2–7.4). Moreover, cancer cells exhibit a pronounced pH gradient across intracellular compartments, ranging from early endosomes (pH 6.0–6.5) to late endosomes (pH 5.0–6.0) and lysosomes (pH 4.0–4.5), largely driven by elevated glycolytic activity under hypoxic conditions [42].

### Effects of 1400W-OPA NPs on NO production

We used LPS-stimulated RAW264.7 macrophages as a classical inflammatory model to assess NO production. First, the cytotoxicity of free 1400W and the NPs (at an equivalent 1400W dose) was evaluated by MTT assay after 24 h of incubation. The NPs showed less cytotoxicity when compared with free 1400W. Indeed, 1400W induced significant cytotoxicity at 40 µM compared with the control (p < 0.01) and this toxicity was increased at 100μM (p < 0.001). Whereas the NPs showed no significant toxicity at 60μM (Figure 2a). This difference is likely attributable to the sustained-release behavior of the NPs, which results in a lower effective concentration of 1400W over 24 h and consequently reduced acute toxicity. Similar trends have been reported for many NP systems *in vitro*. In short-term assays (24–72 h), free drugs are immediately available at high concentrations, leading to rapid cellular exposure and pronounced biological effects, whereas sustained-release NPs deliver the payload gradually, producing lower instantaneous concentrations and weaker acute responses [43, 44]. Nevertheless, compared with free drugs, NPs typically show higher accumulation in tumor and edematous tissues due to the EPR effect [45].

**Figure 2.**
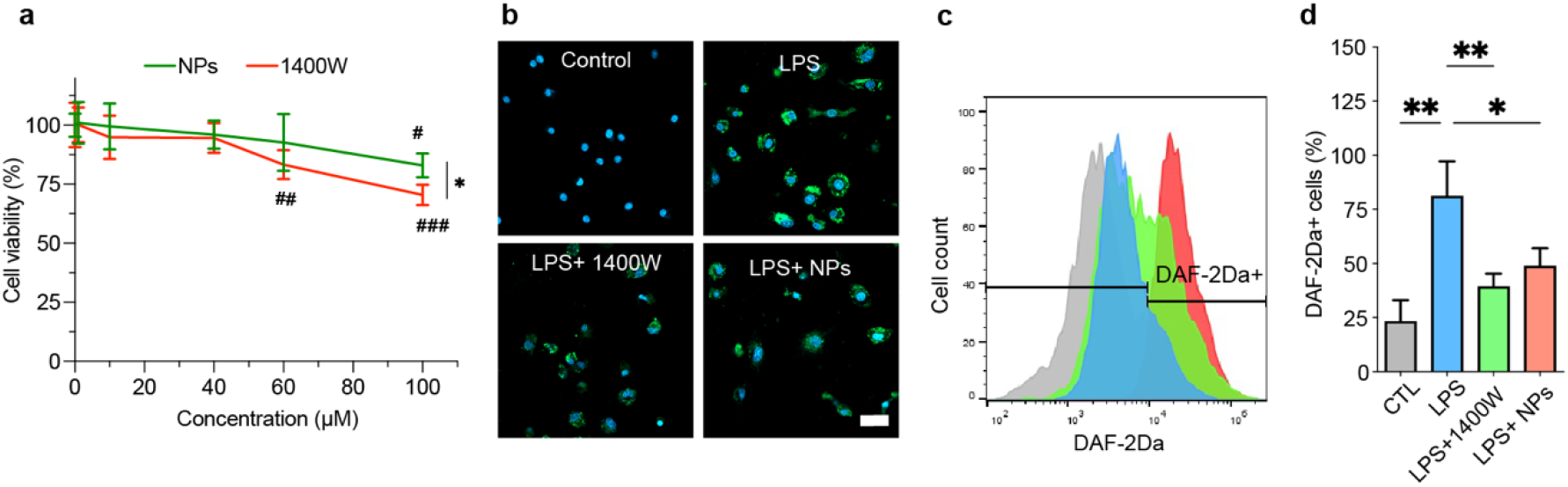
Effects of 1400W and the NPs on LPS-induced NO levels in RAW264.7 cells. **a.** Viability of the cells treated without or with various concentrations of the 1400W and NPs assessed by MTT assay. t-test (at one time point between two treatment) or one-way ANOVA with Tukey’s post hoc test (multiple was used to analyze the data. *p < 0.05 between NPs and 1400W; # p<0.05, ## p<0.05 and ### p<0.05 vs control. Data are expressed as mean ± SD (n=3). **b**. Confocal microscopy images of RAW264.7cells stained with DAF-2Da and incubated with LPS (500 ng/ml), and then without or with 20 μM free 1400W, or the NPs. **c**. Representative flow cytometry histogram of NO levels in DAF-2Da-stained RAW264.7 cells following incubation with the treatments. **d**. Quantification of high-fluorescence DAF-2Da-positive RAW264.7. The comparison was done by one-way ANOVA with Tukey’s post hoc. Data are expressed as mean ± SD (n=3). *p < 0.05, **p < 0.01., and ***p < 0.001. # p<0.05, ## p<0.05 and ### p<0.05 vs. respective control.

Therefore, 20 and 40 µM NPs were selected for subsequent evaluation of effects on NO synthesis, as this concentration did not affect cell viability. Stimulation of RAW264.7 macrophages with 500 ng/mL LPS resulted in a significant increase in NO production, as detected by DAF-2Da fluorescence staining and flow cytometry analysis (p < 0.01 vs. control, Figure 3b–d). This response is consistent with the role of LPS in activating TLR4 in macrophages, initiating intracellular signaling cascades such as NF-κB and MAPK pathways, which regulate iNOS gene transcription and promote NO production [46]. Prolonged NO exposure contributes to cancer progression, particularly by facilitating immune evasion and remodeling of the tumor microenvironment [18]. Compared with LPS-treated cells, treatment with 20 µM free 1400W or an equivalent concentration of NPs significantly reduced NO production, as indicated by decreased DAF-2Da green fluorescence intensity in both confocal microscopy images (Figure 5a) and flow cytometry quantification (Figure 5b, c). Accordingly, RAW264.7 cells treated with either free 1400W or 1400W-OPA NPs exhibited a significant ability to suppress LPS-induced NO synthesis (p < 0.01 and p < 0.05 vs. control, respectively). These findings confirm the capability of NPs to effectively deliver 1400W and inhibit iNOS activity, supporting their potential application in cancer therapy *in vivo*.

**Figure 3.**
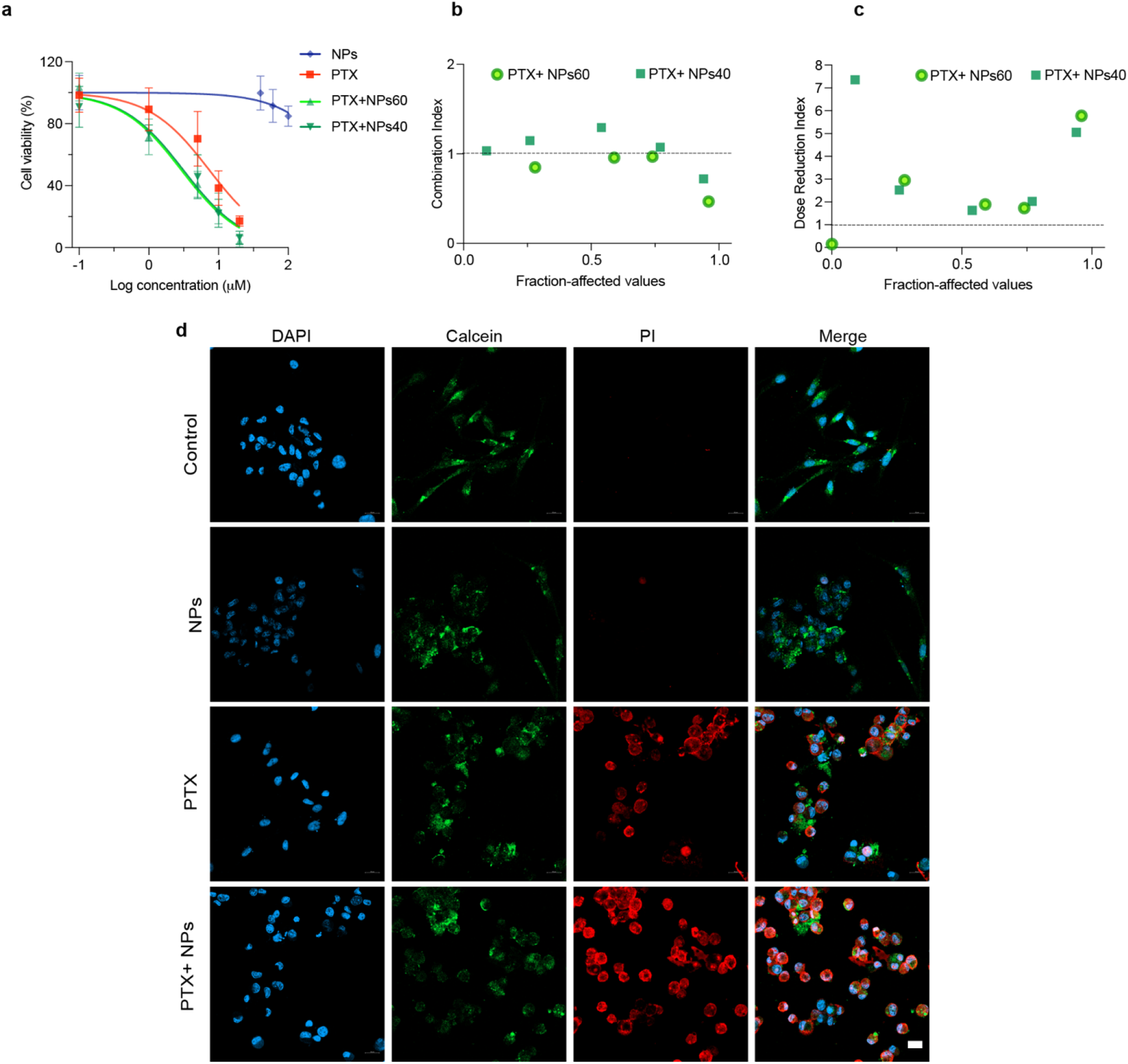
Effect of PTX and NPs on breast cancer cell proliferation. **a.** Cell viability of MDA-MB-231 cells treated in the absence (control) or presence of 20 µM NPs, 1µM PTX, and a combination of NPs with PTX. **b**. Combination index of synergy between PTX and NPs assessed by Chou–Talalay analysis. **c**. Dose reduction index analysis of PTX and NP combinations assessed by Chou–Talalay analysis. **d**. Calcein/PI staining of cells treated with 1 µM PTX and/or 20 µM NPs. Data are expressed as mean ± SD (n = 3). Scale bar shows 20μm.

**Figure 4.**
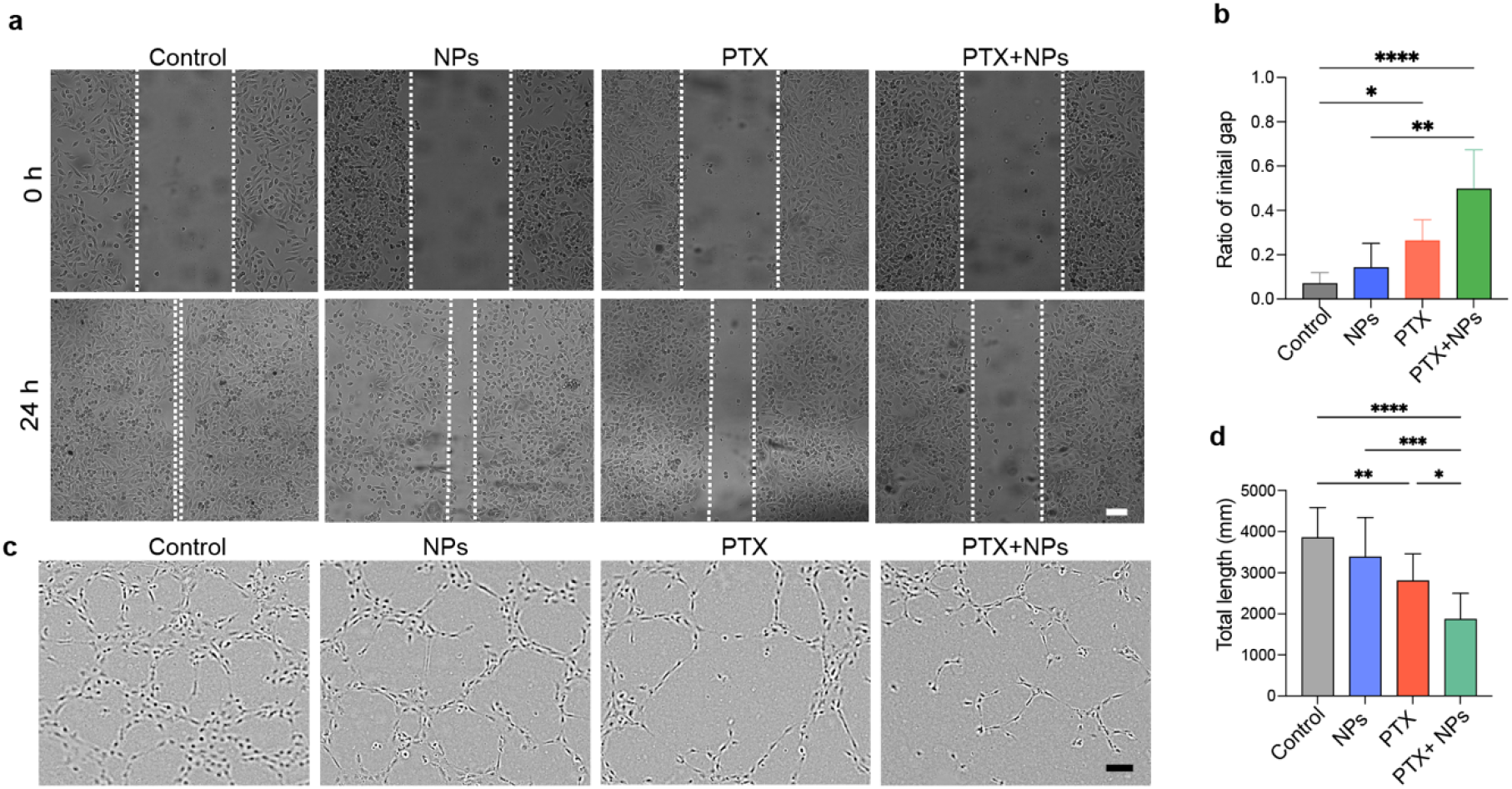
Effects of PTX and nanoparticles (NPs) on breast cancer cell migration and vascular network formation. **a.** Representative images of the scratch wound–healing assay in MDA-MB-231 cells treated in the absence (control) or presence of NPs (20 µM), PTX (1 µM), or their combination. **b**. Quantitative analysis of wound gap closure after 24 h, using Kruskal-Wallis test with Dann’s multiple comparison test. **c**. Representative images of HUVEC tube formation following VEGF stimulation (50 ng/mL) and subsequent treatment with NPs (20 µM) and/or PTX (1 µM). **d**. Quantification of the total tube length formed under each treatment condition. One-way ANOVA with Tukey’s post hoc test was used to evaluate the differences between the groups. Scale bars represent 100 µm. Data are presented as mean ± SD (n = 6). Statistical significance is indicated as *p < 0.05, **p < 0.01, and ****p < 0.0001.

### Synergistic cytotoxic effects of the NPs and PTX

Like what was observed in RAW264.7 cells, the NPs reduced the viability of MDA-MB-231 cells at 100 µM; however, this concentration is not considered sufficiently potent for effective cancer cell killing. Nevertheless, treatment with NPs significantly enhanced the cytotoxicity of PTX. PTX alone exhibited an IC□□ of 7.4 ± 1.7 µM, whereas co-treatment with NPs at 20 and 40 µM reduced the IC□□ to 3.2 ± 0.8 and 2.9 ± 0.6 µM, respectively. Notably, although NPs at these concentrations were not cytotoxic on their own, their combination with PTX resulted in an approximately 2.5-fold reduction in the IC□□ of PTX (Figure 3a).

These interactions were also analyzed using the Chou–Talalay method. The fraction-affected values represent the fraction of cells inhibited following drug exposure [47]. As shown in Figure 3b, several combinations of NPs and PTX yielded CI < 1, indicating synergistic cytotoxicity, with the strongest synergy observed for the combination of PTX with NPs at 40 µM.

The DRI quantifies the extent to which the dose of each agent in a combination can be reduced relative to monotherapy while maintaining the same level of effect. As shown in Figure 3c, the dose of PTX in combination with NPs could be reduced by approximately 3–6-fold to achieve effects comparable to PTX alone. This was further supported by calcein/PI staining, which showed that co-treatment with 20 µM NPs and 1 µM PTX increased the population of apoptotic and necrotic cells, as evidenced by a reduced proportion of calcein-positive cells and increased PI staining (Figure 3d).

The role of NO in cancer biology is complex and multifaceted. It is well established that high NO levels induce nitrosative and oxidative stress, activating apoptotic pathways and promoting cell death, whereas low NO levels are protective and proliferative and can render cells resistant to cytotoxic agents such as chemotherapeutic drugs [4, 48]. The cytotoxic effects of NO have also been shown to synergize with cancer therapies, particularly in studies using NO-donor molecules that generate high NO concentrations as adjuvant treatments [45].

NO plays a complex, context-dependent role in regulating cell migration and metastatic behavior, acting either as a stimulator or suppressor of tumor progression depending on its concentration, duration of exposure, and the cellular context [23]. Increased NO signaling has been linked to enhanced cancer cell motility and invasiveness through modulation of pathways such as PI3K/Akt, MAPK, and mTOR, as well as through NO-driven cytoskeletal remodeling, all of which are essential for metastatic spread [23, 49].

Beyond tumor cell migration, NO functions as a key downstream effector of VEGF signaling, mediating endothelial cell proliferation, survival, and migration through both cGMP-dependent and independent mechanisms, thereby supporting angiogenic processes [50]. Therefore, we investigated the effects of the NPs on cell migration and angiogenesis using a scratch wound– healing assay and tube formation assays respectively. Although the NPs did not significantly affect the closure of the gap or tube formation in HUVECs they significantly increased the effects of PTX. In particular, the gap of the scratched area was 1.9-fold larger in PTX+ NPs when compared with PTX alone. HUVECs tube formation is a well-established model to study angiogenesis *in vitro* [51], and the NPs resulted a significantly potentiated the ability of PTX in inhibition of tube formation when compared with PTX alone (p < 0.05). These data corroborated the findings of cell toxicity in MDA-MB-231, hence it is expected that at least part of the effects prevent proliferative effects of NO generated by iNOS.

Pharmacological suppression of NO production has been shown to mitigate pro-tumorigenic signaling and enhance the efficacy of anticancer therapies [18, 48, 52]. Granados et al. reported that combined treatment with NG-monomethyl-L-arginine (L-NMMA) and docetaxel significantly inhibited tumor growth in an orthotopic MDA-MB-231 model [52]. However, because L-NMMA functions as a non-selective NOS inhibitor, concomitant inhibition of eNOS can result in adverse hemodynamic effects, including hypertension and reduced cardiac output. To mitigate these effects, amlodipine was incorporated to maintain cardiovascular stability. This combinatorial approach, integrating non-selective NOS inhibition with docetaxel, led to improved survival outcomes compared with docetaxel monotherapy. Notably, L-NMMA has progressed to phase I and II clinical trials as a chemosensitizing agent for treatment-resistant TNBC [53].

In a xenograft model of adenoid cystic carcinoma of the oral floor, administration of the highly selective iNOS inhibitor 1400W, either alone or in combination with a CXCR4 antagonist, resulted in marked suppression of tumor growth [54]. Importantly, 1400W demonstrated significantly greater efficacy than L-NAME in reducing subcutaneous tumor burden, which was attributed to elevated iNOS expression and the limited iNOS selectivity of L-NAME. Beyond tumor growth inhibition, treatment with 1400W induced apoptosis and substantially attenuated tumor-associated angiogenesis and cellular proliferation [54]. Despite these promising biological effects, the translational utility of 1400W, as with many small-molecule inhibitors, is limited by rapid systemic clearance and inadequate tumor accumulation. Accordingly, nanocarrier-based delivery systems, such as the NPs developed in the present study, represent a promising strategy to improve 1400W delivery and exploit the EPR effect, thereby addressing key pharmacokinetic limitations associated with 1400W and iNOS inhibitors more broadly.

## Conclusion

In this study, we developed stable nanosized formulations via Schiff base conjugation between OPA and 1400W to enable controlled inhibition of iNOS. The resulting NPs exhibited pH-responsive drug release behavior and effectively suppressed NO production in macrophages, demonstrating their functional activity. Although the NPs did not induce cytotoxicity as monotherapy, they significantly enhanced the anticancer efficacy of PTX in TNBC cells. Specifically, the combination treatment synergistically increased PTX-induced cytotoxicity and potentiated its inhibitory effects on cancer cell migration as well as on endothelial tube formation in HUVECs, indicating a potential dual impact on tumor cell aggressiveness and angiogenic processes.

The absence of *in vivo* evaluation represents a limitation to our study considering it precludes assessment of key pharmacokinetic parameters, including circulation half-life, tumor accumulation, and biodistribution, as well as validation of the therapeutic efficacy of the NPs in combination with PTX in tumor-bearing models. Therefore, future studies incorporating appropriate *in vivo* models will be essential to determine whether the enhanced biological activity observed *in vitro* translates into improved antitumor outcomes and to fully establish the therapeutic potential of this nanocarrier-based strategy.

## Acknowledgements

The authors would like to thank Ms. Françoise Gregoire and Ms. Nargis Bolaky for their administrative and technical assistance with this project. This project receives funding from the European Union’s Horizon 2020 research and innovation programme under the Marie Skłodowska-Curie Grant Agreement No 101034324, as well as a grant from the Jaumotte-Demoulin Foundation. Some of the equipment used in this study was financed, in whole or in part, by the Walloon Region through the Technology Platforms of Excellence: ‘Alternative to animal experimentation’ and ‘Biogreen’. The authors gratefully thank the Centre d’Instrumentation en Resonance Magnétique – CIREM (Université libre de Bruxelles – ULB, Belgium) for providing support and access to its infrastructure, in particular the NMR spectrometer J400 funded by the Fonds de la Recherche Scientifique (F.R.S.-FNRS – GEQ2011-2.5014.12) and the Fonds d’Encouragement à la Recherche (FER-ULB). The Center for Microscopy and Molecular Imaging (CMMI) is supported by grant 411132-957270 from the European Regional Development Fund and the Walloon Region (Wallonia-Biomed; project “CMMI-ULB”). The authors thank Louise Conrard (CMI) for her support with TEM analysis. MA benefited from Grant n° 72229 from Tabriz University of Medical Sciences to support her Ph.D. thesis.

